# A waveguide imaging platform for live-cell TIRF imaging of neurons over large fields of view

**DOI:** 10.1101/2019.12.13.874545

**Authors:** Ida S. Opstad, Florian Ströhl, Marcus Fantham, Colin Hockings, Oliver Vanderpoorten, Francesca W. van Tartwijk, Julie Qiaojin Lin, Jean-Claude Tinguely, Firehun T. Dullo, Gabriele S. Kaminski-Schierle, Balpreet S. Ahluwalia, Clemens F. Kaminski

**Author notes:** Both authors contributed equally to this work (Ida S. Opstad, Florian Ströhl).

## Abstract

Large fields of view (FOVs) in total internal reflection fluorescence microscopy (TIRFM) via waveguides have been shown to be highly beneficial for single molecule localisation microscopy on fixed cells [1, 2] and have also been demonstrated for short-term live-imaging of robust cell types [3–5], but not yet for delicate primary neurons nor over extended periods of time. Here, we present a waveguide-based TIRFM set-up for live-cell imaging of demanding samples. Using the developed microscope, referred to as *the ChipScope*, we demonstrate successful culturing and imaging of fibroblasts, primary rat hippocampal neurons and axons of *Xenopus* retinal ganglion cells (RGC). The high contrast and gentle illumination mode provided by TIRFM coupled with the exceptionally large excitation areas and superior illumination homogeneity offered by photonic waveguides have potential for a wide application span in neuroscience applications.

---

TIRFM provides an effective means for the spatially confined illumination of a sample close to the coverslip/substrate via evanescent fields [6–8]. It provides particular advantages for fluorescence imaging as out of focus signal is intrinsically avoided leading to high signal to noise ratios and image contrast. In addition to the molecular specificity afforded by fluorescence imaging and image contrast, TIRFM reduces the overall illumination dose on the sample. This minimises phototoxicity, making TIRF the method of choice for many live-cell imaging applications with delicate samples such as live neurons [9].

TIRFM is usually accomplished by using a large numerical aperture (NA) objective lens for both the excitation and detection paths. Unfortunately, the high magnification of lenses required for TIRFM limits the accomplishable FOVs and thus imaging throughput. This makes conventional TIRFM impractical for extended and often fast moving samples such as neurons and their organelles [10]. This FOV restriction is removed if waveguides are used for TIRF illumination and in principle, arbitrarily large areas could be achieved through appropriately designed waveguide geometries (width and length). Because the excitation and detection paths are completely decoupled from one another, full flexibility in choice of the imaging objective lens is retained, allowing for control over the FOV size, as illustrated on fibroblasts in Suppl. Figure S1. In the so called *ChipScope* microscopy system [1], multiple colours can be admitted simultaneously into the photonic chip, enabling the simultaneous TIRF excitation of multiple fluorophores (see Suppl. Figure S2).

Waveguides have previously been shown to be a viable growth substrate for cell culture [3, 4], but to fully exploit the gentle TIRF illumination for live-cell image applications, especially in the neurosciences, additional considerations and adaptations must be made to maintain the cells alive under suitable conditions. The scope of this work was to adapt a waveguide TIRF microscopy set-up for the imaging of sensitive cell types like primary neurons, and to develop means of performing measurements on primary cell-cultures on photonic chips. These are demanding and challenging cells to grow in general and especially on waveguide materials (illustrated in Suppl. Figure S3), as the surface properties are different compared to cover glasses, which are currently standard for neuronal culturing.

Cultured neurons from *Xenopus laevis* (African clawed frog) are viable at room temperature under atmospheric levels of oxygen and CO_2_, making them an attractive and practical choice for studies requiring prolonged live imaging and a suitable initial test specimen for the *ChipScope*. The axons from *X. laevis* RGC are for example a well-established model system for the study of axon guidance via TIRFM [11]. The growth cones at the tip of developing axon terminals are flat, hand-shaped structures sensitive to extracellular chemical and molecular stimuli and support axon navigation during embryonic development [12]. To image the growth of live RGC axons in culture, we explored the capabilities of our waveguide imaging platform in simultaneously capturing tens of growth cones of far-reaching axons from explanted *Xenopus* eye primordia. Different from previous waveguide imaging implementations, we employed water dipping objective lenses, which greatly facilitate high resolution live-cell imaging and permit access to the sample during imaging, e.g. to optimise labelling conditions or study the response of different treatments *in actu* [13]. Live-cell imaging results of filamentous actin in developing axons and growth cones of RGC are shown in Figure 1. The benefit of TIRFM over episcopic (EPI) illumination is apparent when comparing identifiable single cortical filaments of growth cones, as shown in Figure 1, panels d-f. Additionally, the vastly increased FOV supports quantitative analyses.

**Figure 1:**
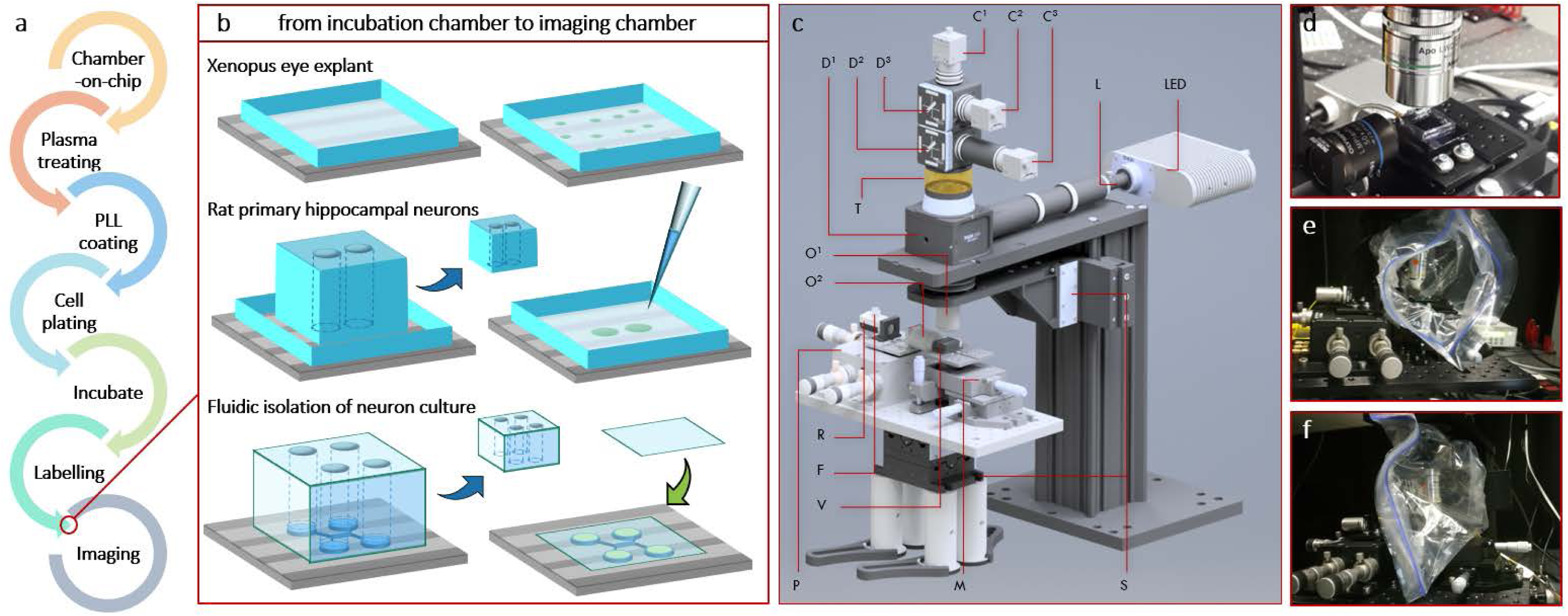
Chip microscopy is a method for imaging of large areas of live Xenopus RGC axons in TIRF. All panels show images of the same living *Xenopus* eye explant cultured on chip and labelled with SiR-Actin. a) Overview image captured using 0.3NA water dipping objective in both TIRF (upper panel) and BF (lower panel) mode. b) Overview image captured using 1.1NA water dipping objective using EPI (left panel) and TIRF (right panel) illumination modes. d-f) Excerpts from region indicated in b, comparing available growth cone details for different NAs in TIRF and EPI illumination modes. TIRF illumination together with 1.1NA in panel f reveals the most identifiable single cortical filaments. TIRF images were obtained via summation of 100 frames illuminated by different waveguide illumination modes. Scale bars: a) 100 µm, b) 50 µm, d-f) 10 µm.

While *Xenopus* neurons can be imaged under ambient conditions, mammalian cells require 37°C and 5% CO_2_. To allow for long term imaging of mammalian neurons under physiologically relevant conditions, we equipped the *ChipScope* with a heater system and a custom-made environmental chamber. As laser coupling together with its delicate piezo stage electronics precludes the use of common commercially available microscope stage incubators, we custom designed a chamber connected to a commercial (Okolab) stage top incubator system, suppling 5% CO_2_ and high humidity. Our chamber is of transparent, flexible and low heat conductivity low-density polyethylene (LDPE) thermoplastic, that can be easily cut and stretched to tightly fit around all necessary microscope components, while maintaining easy access for changing sample or objective through a zip open/close mechanism (Figure 2e-f). A heating strip and temperature sensor are fitted to the sample holder to maintain the chip at 37°C during imaging experiments. Further details are provided in Suppl. Note 1.

**Figure 2:**
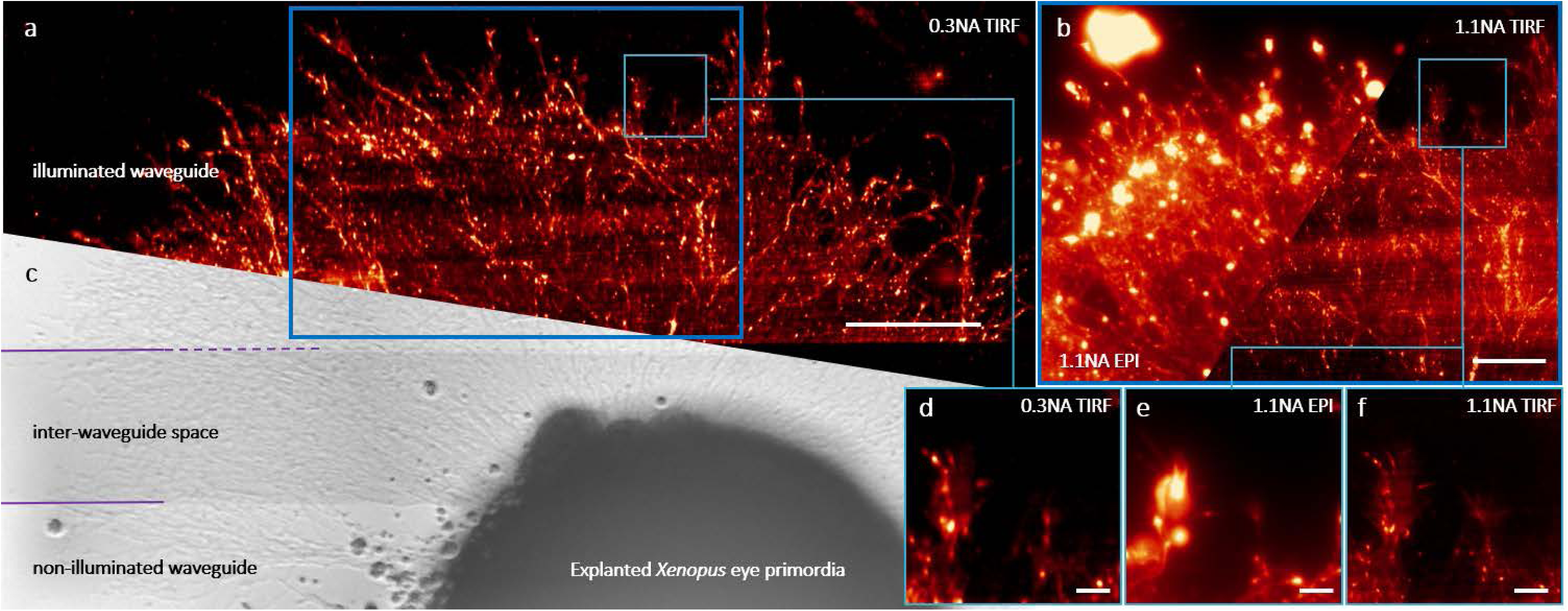
Chip microscopy can be adapted for live-cell imaging and various cell culture approaches. a) Experimental flowchart. b) Photonic chip preparation for cell culture and imaging. *Top*: chambers for *Xenopus* retinal ganglion cells, *middle:* chambers for rat hippocampal neurons, *bottom:* two layers of PDMS for cultivating neurons in microgrooves. Thin bottom layer remains for imaging with a coverslip to reduce evaporation. c) *ChipScope* model. This upright microscope enables TIRF, EPI and BF imaging with up to three colours simultaneously. A detailed description of the optical set-up is provided in Suppl. Note 1. d) Microscope stage with waveguide chip and imaging chamber prepared for use with water dipping objective. The horizontal objective is for laser coupling into the waveguides. e) ChipScope with open incubation chamber for easy access to sample and objective. f) Closed incubator supplied with high humidity and 5% CO_2_ from a conventional stage top incubator.

Standard protocols for culturing rat hippocampal neurons require tall, slim chambers (as in Figure 2b, second row). These culture chambers are incompatible with upright microscopes featuring short working distance, high NA objectives, while inverted microscopy is rendered impractical by the opaque base layer of silicon that forms a supporting platform for the waveguides. We solved this difficulty via separate custom-made wells for culturing and imaging, as displayed in Figure 2b. After growing, the neurons on-chip in their preferred polydimethylsiloxane (PDMS) cell culture wells, we removed the tall PDMS blocks right before imaging in favour of wider and lower ‘fences’ adapted for the particular waveguide chip and imaging objective of choice. To monitor the on-chip hippocampal cultures in the time between excision and laser lab TIRF imaging, we built a simple upright microscope that could conveniently be fitted on a standard biological lab bench, see Suppl. Fig S4. A flowchart of our on-chip sample preparation is shown in Figure 2a-b and the imaging set-up in Figure 2c-f.

The satisfactory performance of this incubator was validated through longer term live-cell imaging of delicate primary hippocampal neurons (excised from rat embryos). The results, displayed in Figure 3, show the microtubule network imaged in TIRF, EPI and brightfield (BF) mode (from left to right). After 1.5h of imaging, the primary neurons were observed to be in healthy condition (Suppl. Fig. S5). To the best of our knowledge, this is the first chip-based imaging system with incubation chamber, successfully adapted for live-cell imaging of mammalian neurons.

**Figure 3:**
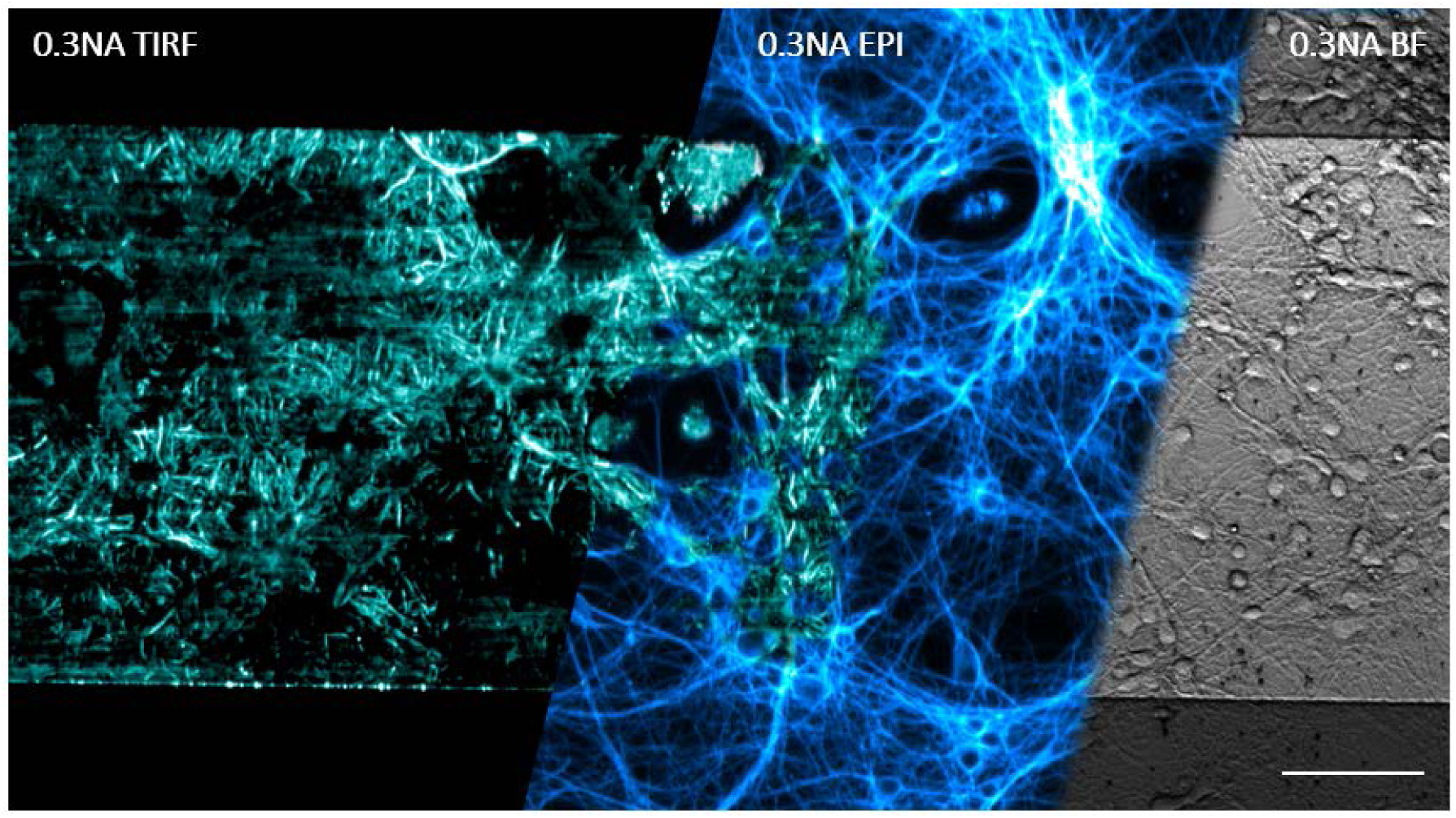
Rat hippocampal neurons can be cultured on chip for weeks and imaged live for hours in ChipScope incubator. All panels show live-cell imaging of neurons labelled with SiR-Tubulin on adjacent regions of the same waveguide. *Left*: TIRF image obtained by mode-averaging as in Figure 1. *Middle*: Single plane EPI image overlaid with part of the corresponding TIRF image. *Right*: BF image. Scale bar: 100 µm.

Advanced cell culture approaches are becoming more and more important in neuroscience research, such as microgroove cell culture chambers used in studies of axonal injury or protein transport [14–16]. We therefore sought to use PDMS microgroove chambers in combination with the photonic chip imaging system. One complication is that the cell culture chambers are usually permanently bonded to the substrate, but the chips are – at present – too expensive to be disposed of after a single experiment. We found that clean PDMS was sufficiently tacky to adhere to the photonic chip and contain media while the neurons grow. Secondly, as the chips are opaque, signal must be collected from above. This requires the PDMS to be thin enough to match the working distance of the desired imaging objective, and to be uniform and optically clear, so that high quality imaging can be performed through the PDMS layer. However, such a thin PDMS layer would not contain sufficient media to support the neurons and overcome evaporation. We addressed these challenges by using a second PDMS layer on top of the microfluidic devices to contain the media, which was removed before imaging, as depicted in Figure 2b, bottom row. As displayed in Figure 4, we successfully performed TIRF, EPI and BF microscopy on photonic chips through PDMS microgrooves of living primary hippocampal neurons using a high NA silicon oil immersion objective. The results show that this multimodal imaging scheme is feasible, although the TIRF imaging success in this case was modest as the axons appeared to have detached from the waveguide TIRF excitation area. This might be addressed in future experiments e.g. by selective coating of the waveguide surface with poly-l-lysin, without coating the microchannel walls, thus removing possible attachment points for the growing axons above the waveguide surface. Another challenge with fluorescence imaging in microgrooves concerns the labelling and consecutive washing steps required in many protocols, which are difficult to achieve in the narrow channels. In our data, this resulted in high background signal by the used MitoTracker label, causing the mitochondria to be barely visible in the EPI image of Figure 4e. Supplementary Note 4 provides details on the procedures and microgroove production steps.

**Figure 4:**
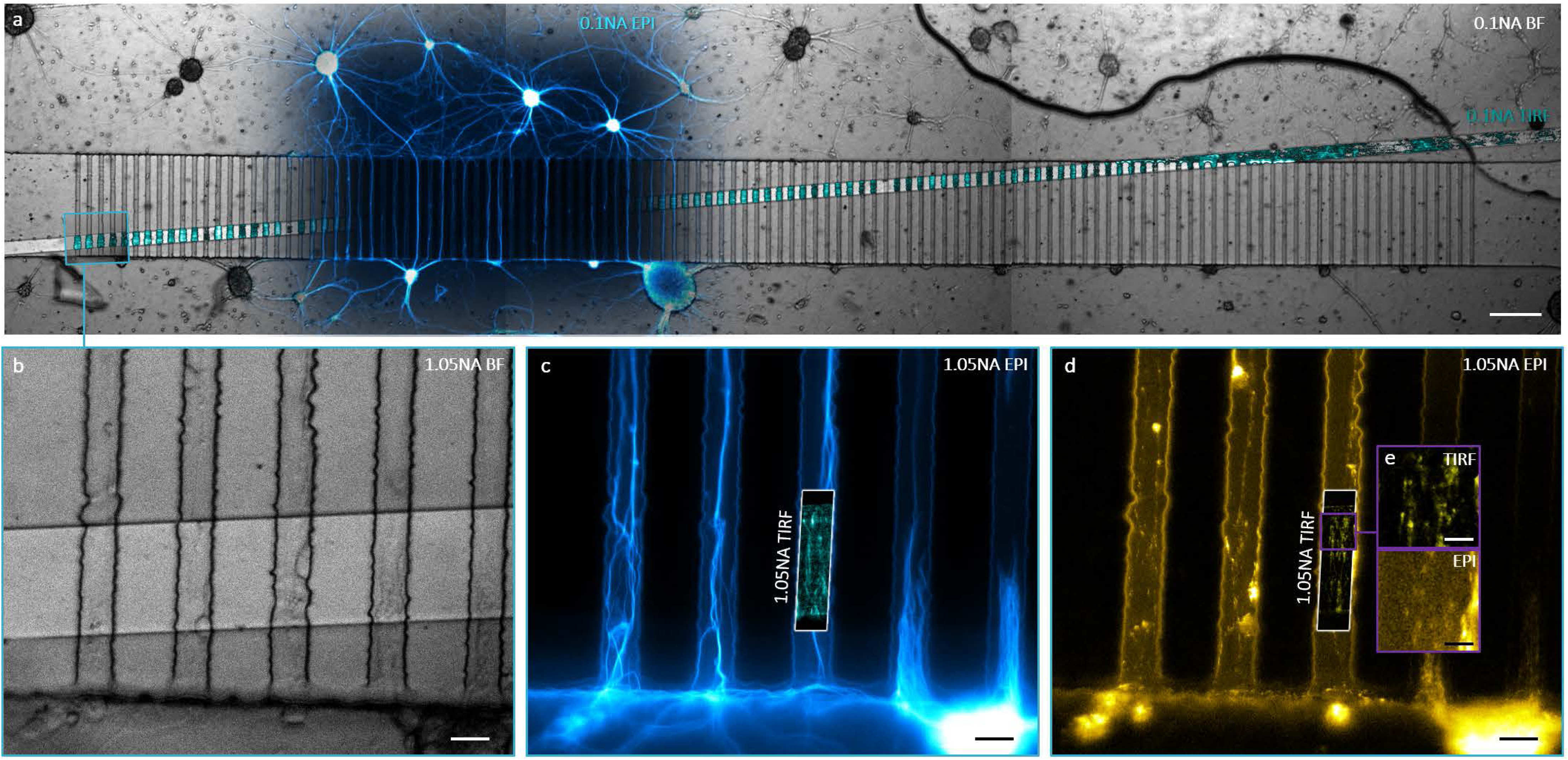
Advanced cell culture approaches like microchannel devices can be combined with large FOV chip TIRFM. All panels show rat hippocampal neurons cultured in a chip-microchannel combi device and labelled using SiR-Tubulin and MitoTracker Orange. a) Stitched overview image acquired using a 4X 0.3NA air objective. BF mode overlaid with EPI (blue) and TIRF (cyan) illumination. Scale bar: 200 µm. b)-d) Images acquired using a 30X 1.05NA silicon oil immersion objective, scale bars are 20 µm. b) BF image of microchannels on waveguide. c) Corresponding EPI image of microtubules with TIRF image inlay. d) EPI image of MitoTracker Orange with TIRF inlay. e) Zoomed view of TIRF and EPI image of mitochondria in microchannels, scalebar 5 µm.

In summary, we have adapted photonic chip large area TIRFM for live-cell neuroimaging applications by developing on-chip cell culture protocols and integrating chip microscopy with a heater system and incubation chamber. We have shown successful cell culture of primary rat hippocampal neurons and excised Xenopus eye primordia and performed large FOV live-cell TIRFM of these delicate cell types. We have demonstrated imaging on combined microgroove-waveguide devices and the capabilities of our system. We expect the integration of environmental control with the unique advantages of TIRF illumination provided by waveguides to inspire and enable many new imaging applications for photonic chips.

## Additional information

### Author contributions

ISO led the project and performed the imaging. FS designed the ChipScope. ISO and FS built and calibrated the system. BSA, FTD and JCT provided waveguides and advised on chip microscopy. ISO, FS and OV designed and installed the custom imaging incubator. MF wrote the ChipScope control software. CH excised, cultured and prepared labelled samples of primary rat hippocampal neurons on chip. CH advised on the incubator and helped plan and conduct the imaging experiments. OV designed and fabricated the microchannel moulds. ISO, CH and OV made microchannels and PDMS chambers. JQL and FWT provided expertise on *Xenopus* biology and handling. FWT performed *Xenopus* excisions and on-chip culturing and advised on the labelling and imaging of RGCs. CFK and BSA provided supervision and funding for the project. GSKS provided facilities for primary neurons excision and cell culture, and CFK microscopy lab area and equipment. ISO and FS wrote the manuscript. All authors contributed to or commented on the manuscript.

## Acknowledgements

The authors thank Omid Siddiqui for performing simulations on heat transfer of the sample stage and Christine E. Holt for providing means of *Xenopus* culturing and loan of silicon oil objectives. This work was supported by grants from the European Research Council, project number 336716 to BA and the Tematiske Satsinger funding program to BA by UiT The Arctic University of Norway. FS acknowledges funding via Horizon 2020 Marie Skłodowska-Curie actions (#836355). CFK acknowledges funding from the Physical Sciences Research Council (EP/H018301/1), Medical Research Council (MR/K015850/1 and MR/K02292X/1), Wellcome Trust (089703/Z/09/Z), MedImmune | AstraZeneca and Infinitus China Ltd.

## Supplementary information

### Suppl. Note 1: Optical set-up

For illumination and imaging of the photonic waveguide chips, a custom upright microscope was used, whose design was adapted from the one used in a previous publication [1]. A 3D model of the microscope is displayed in Figure 2c of the main article, highlighting the main components that comprise the frame and stage, chip-TIRF and episcopic illumination, as well as detection modules.

The frame was split in two parts, with the microscope body being from a modular commercial system (CERNA, Thorlabs) and the stage assembly housing the chip. The chip itself rests on a vacuum stage (HWV001) and is held in place with low vacuum. A three-axis long travel block (RB13M/M, Thorlabs) allows coarse alignment of the chip with respect to the differential micrometre stage and lateral translation of the whole assembly is realised via a motorized two-axis translation stage (PLS-XY, Thorlabs). Focusing onto the sample is performed by translating the imaging objective via a 1-inch travel module (ZFM2020, Thorlabs) with micrometre precision.

Evanescent chip-TIRF illumination of the sample is achieved by focusing laser light into the waveguide chip inputs. The laser light is provided by up to four laser lines at 445 nm with 75mW (OBIS 445LS, Coherent), 488 nm with 150 mW (OBIS 488LS, Coherent), 561 nm with 150 mW (OBIS 561LS, Coherent), and 647 nm with 120 mW (OBIS 647LX, Coherent). The lasers are custom combined using dichroic mirrors (ZT458rdc, ZT514rdc, ZT594rdc, all Chroma) and coupled into a single-mode fibre (S405-XP-custom, Thorlabs) using a commercial fibre coupler (PAF2-A4A, Thorlabs). The fibre output is successively collimated using a reflective collimator (RC04FC-P01, Thorlabs) to minimise chromatic off-set on the coupling facet and the collimated light then focused with a 50x 0.5NA objective (LMPLFLN-BD 50X, Olympus) onto the inputs of the waveguide chips.

For aiding coupling, ease-of-use, as well as access to mode-averaging [2], the reflective collimator and the coupling objective are installed on a differential micrometre stage with piezo drive (MDT630B/M with MAX302/M, Thorlabs). The episcopic illumination light is produced in a four LED combiner (LED4D245, Thorlabs) and delivered to the main frame via a liquid light guide of about 3 mm diameter (LLG0338-4, Thorlabs). It offers illumination with up to four LEDs simultaneously with wavelengths centred on 385 nm, 490 nm, 565 nm, and 625 nm. To counter the broad emission of the 565 nm LED, a bandpass filter (#86-986, Edmund Optics) is custom-installed in the LED combiner. The light guide couples to a set of conditioning optics (WFA2001, Thorlabs) that redirects the light via reflection off a quad-band dichroic mirror (Di03-R405/488/561/635-t3, Semrock) onto the sample through one of the imaging objectives (PLN4XP, Olympus; UPLSAPO30XS, Olympus; CFI Plan Fluor 10X, Nikon; CFI75 Apochromat 25XC W, Nikon).

Produced fluorescence, scattered evanescent light, and back-reflected brightfield light is captured by the respective imaging objective and focused by a tube lens (TTL180-A, Thorlabs, for Olympus objectives and TTL200-A, Thorlabs for Nikon objectives) onto up to three CMOS cameras (UI-3080CP-M-GL Rev.2, IDS), which feature a 3.45 µm pixel size. Detection light splitting is realised by dichroic mirrors (Di03-R561-t1, Di03-R635-t1, both Semrock), and the successive emission filters (FF01-525/45, FF01-600/52, FF01-680/42, all Semrock) are housed by commercial filter cube holders (DFM1/M, Thorlabs) just before the cameras.

The custom incubator was made from a commercially available food bag (item code: 7186606, Sainsbury's Food Bags, Medium Zip & Seal 27.5×26cm x20, Sainsbury’s, UK) and modified to provide openings for light delivery, detection, stage, and environmental control. Environmental control was achieved by coupling a conventional CO_2_ chamber (Okolab) via airtight tubing to the custom chamber. CO_2_ regulation was set to 5%. For heating of the stage, a heat strip with temperature sensor was attached to the sample stage and connected to a temperature controller (TC200, Thorlabs).

### Suppl. Note 2: Labview Software

The ChipScope uses LabVIEW to automate common acquisition tasks and provide a clear user interface, shown in Supplementary Figure S6a. The software controls laser and LED power and switching, stage XYZ position, piezo XYZ position, and camera acquisition. Users can set up a multi-frame acquisition to capture a time-series, a z-stack, a mosaic grid, multi-colour images, or a combination of all these options. The interface is designed to be accessible to new users, minimising training time and thereby increasing the throughput of the microscope.

The block diagram powering the software, shown in Supplementary Figure S6b, has been designed following best-practice LabVIEW paradigms. The block diagram follows the state machine design paradigm, with user interface interactions handled by a separate loop to hardware interface tasks.

This prevents blocking of the user interface front panel, allowing tasks to be interrupted – for example, an acquisition task can be cancelled if the user realises incorrect parameters have been entered.

In addition to the state machine, a producer-consumer design pattern has been implemented for asynchronous image saving. Images acquired from the cameras are added to a memory queue which is read in a separate saving loop. This allows the cameras to acquire images as fast as possible without delay caused by writing the images to disk.

Colour-coded subVIs have been used extensively to ensure the program fits on a single computer monitor and does not become a tangled web of wires. This makes program flow clear to new developers of the software, making it easy to quickly implement new features. The software is provided as an open-source package at https://github.com/laseranalyticsgroup/chip-scope, and can be modified by other researchers for use with their own microscope, or used as a template for creating best-practice LabVIEW programs.

### Suppl. Note 3: Waveguide chip fabrication

Thin waveguides made of high refractive index contrast materials such as silicon nitride (Si_3_N_4_) or tantalum pentoxide (Ta_2_O_5_) on silicon dioxide (SiO_2_) have been demonstrated to provide high evanescent field strength for bioimaging applications [2–4]. For availability reasons, two different waveguide types were used for this work, both based on silicon wafers with a substrate of 2 µm deep pre-oxidised SiO_2_. Si_3_N_4_ waveguides were used for experiments shown in Figures 1, 3 and all Supplementary Figures, and Ta_2_O_5_ for the results displayed in Figure 4.

#### Si_3_N_4_

150 nm Si_3_N_4_ were deposited on the substrate through low-pressure chemical vapour deposition at 800°C. Standard photolithography defined the waveguide geometry using photoresist, and reactive ion etching (RIE) was used to etch the entire Si_3_N_4_ around the designed structures. The remaining photoresist was removed, and a 1.5-2 µm thick top cladding layer was deposited by plasma-enhanced chemical vapor deposition at 300°C. To seed the cells at specific imaging areas, the top cladding was removed using RIE and wet etching as in [5].

#### Ta_2_O_5_

220 nm of Ta_2_O_5_ were magnetron sputtered on the substrate. Parameters used for the deposition were 200°C substrate temperature, 300 W magnetron power, and oxygen and argon flow rates of 5 and 20 sccm, respectively. Conventional photolithography was used to pattern the waveguide structures on a photoresist layer, and argon ion-beam milling removed the entirety of the Ta_2_O_5_ layer around the straight waveguide channels. For the ion-beam milling process, an angle of 45° of the sample considerably reduced the side-wall roughness of the waveguides as compared to conventional flat/0° orientation [6]. After solvent rinsing, the samples were treated with plasma-ashing for ca. 10 min to remove any remaining surface photoresist.

The final wafers were scribed and broken down to ca. 2×3 cm before polishing for smooth edges.

### Suppl. Note 4: Microchannels fabrication

#### Microfluidic master fabrication

Conventional mask UV-lithography was used to fabricate master moulds for thin film PDMS neural cell culture devices compatible with waveguide chips. The chip design consists of two reservoirs with a height of 50 µm to provide two independent chambers for cell cultures and an array of 5 µm high microchannels connecting these. First a 3-inch silicon wafer was spin coated (Spin coater model WS-650MZ-23NPP, Laurell) with SU-8 photoresist (Microchem, 3005 series) at 3000 rpm and prebaked at 95°C for 3 min. The microchannel patterns were transferred into the photoresist via UV-lithography using a film mask (Micro Lithography Services) exposed using the UV-LED setup mentioned in [7]. After post-baking for 3 min at 95°C, the wafer was developed in PGMEA (Propylene glycol monomethyl ether acetate, Sigma Aldrich) and rinsed in isopropanol (Sigma Aldrich) to remove any residues, then dried using a nitrogen gun. Next, a 50 µm thick photoresist layer was spin coated using SU-8 (Microchem, 3050 series) at 3000 rpm and prebaked for 15 min at 95°C. Additional alignment marks in the mask design enabled alignment of the reservoirs film mask with the microchannels. After a second UV-exposure and post baking at 95°C for 4 min, the wafers were developed in PGMEA and isopropanol prior to soft lithography.

### Suppl. Note 5: Xenopus laevis embryos and retinal cultures

#### Ethics statement

All animal experiments were approved by the University of Cambridge Ethical Review Committee in compliance with the University of Cambridge Animal Welfare Policy. This research has been regulated under the Animals (Scientific Procedures) Act 1986 Amendment Regulations 2012 following ethical review by the University of Cambridge Animal Welfare and Ethical Review Body (AWERB).

#### Xenopus laevis embryos

Xenopus laevis eggs were fertilized in vitro and embryos were raised in 0.1X Modified Barth’s Saline (MBS; 8.8 mM NaCl, 0.1 mM KCl, 0.24 mM NaHCO_3_, 0.1 mM HEPES, 82 μM MgSO_4_, 33 μM Ca(NO_3_)_2_, 41 μM CaCl_2_) at 14-20°C and staged according to the tables of Nieuwkoop and Faber [8].

#### Primary Xenopus retinal cultures

A PDMS chamber capable of holding 500 μL of solution was fitted onto a clean and dry photonic chip. The chip-chamber device was plasma treated and poly-l-lysin coated as described under Suppl. Note 6, before 1-hour incubation with 10 μg/ml Laminin (Merck) in phenol red-free Leibovitz’s L-15 medium (ThermoFisher). The chips were then washed once with culture medium (60% L-15, 40% ddH_2_O, 1X antibiotic-antimycotic (ThermoFisher), pH 7.6-7.8) before filling the PDMS chambers with 500 μL of culture medium.

To obtain retinal ganglion cell cultures, Xenopus laevis embryos at stage 33-36 were first washed three times in embryo wash solution (0.1X MBS, 1X antibiotic-antimycotic, pH 7.6-7.8), then anaesthetised using a solution of tricaine methanesulfonate (MS-222) (Merck) in 1X MBS with 1X antibiotic-antimycotic (pH 7.6-7.8). Subsequently, embryos were transferred to a dissection dish containing equal volume of MS-222 solution and culture medium. Eye primordia were dissected (covering skin was cut open and surrounding tissue carefully severed using holder-mounted 0.10-mm Minutien Pins (Fine Science Tools)). Eye primordia were subsequently washed three times in culture medium and then transferred onto the chip. Primordia were positioned onto waveguides. After 24 hours, cultures were taken for imaging.

While aligning the chips for optimized laser coupling, the cultures were incubated with 0.5 µM SiR-Actin (Spirochrome Ltd., Stein am Rhein, Switzerland) and 5 µM verapamil (Spirochrome Ltd.) for about an hour, before carefully replacing the staining solution with room temperature culture medium for imaging.

### Suppl. Note 6: Hippocampal neurons, excision and cultivation

#### Ethics statement

All animal work conformed to guidelines of animal husbandry as provided by the UK Home Office. Animals were bred and supplied by Charles River UK Ltd. (Margate, UK), Scientific, Breeding and Supplying Establishment, registered under Animals (Scientific Procedures) Act 1986, and AAALAC International accredited, and sacrificed under schedule 1; procedures that do not require specific Home Office approval.

#### PDMS cell culture chambers

Cell culture chambers were made of PDMS (polydimethyl siloxane, Sylgard^®^ 184 Silicone Elastomer, Dow Corning, Midland, MI, US) mixed 1:10 (curing agent:base), degassed, and cured at 65°C for 1 hour. The microgroove cell culture devices used in this study were based on designs from Taylor and colleagues [9], with taller microgrooves (5 µm). As shown in Figure 2b, a thin (<1 mm) layer of PDMS was cast on an SU-8 (Microchem) master to make the microgroove chamber, and a second thick (~5 mm) layer was cast to sit on top and hold sufficient media for culturing. Inlets were cut with a 6 mm biopsy punch, and the devices were cleaned of dust using Scotch Magic tape (3M, UK). Simple well chambers were produced from flat ~5 mm thick PDMS with wells cut using a 6 mm biopsy punch. Thin layer PDMS devices were attached onto waveguide chips using their natural adherence and put into an oven for 30 min at 65°C to improve attachment. Afterwards, devices were plasma treated with a plasma cleaner (Model Zepto, Diener Electronics) for 40s at 40W in a 0.35 mbar oxygen atmosphere to render the inside of the microfluidic chip hydrophilic. Immediately after plasma treatment, surfaces were coated with 0.1 % poly-l-lysine (Sigma-Aldrich, Darmstadt, Germany) for 1 hour, under UV light to sterilize the devices. Chambers were washed with water and dried before loading cells.

#### Primary rat embryonic hippocampal neuron culture

Sprague-Dawley E18 pregnant rats were sacrificed and embryos were dissected in HBSS (Gibco, Waltham, MA, US) with Antibiotic-Antimycotic (Gibco) to extract the hippocampus. Hippocampi were incubated in 0.1% trypsin (Worthington Biochemical Corporation, Lakewood, US) and 0.05% DNase (Sigma Aldrich) in DMEM (Gibco) for 20 min, washed four times with 0.05% DNase in DMEM, and then dissociated by trituration. Cells were pelleted by centrifugation and resuspended in DMEM with 10% FBS (Gibco). 2×10^4^ cells loaded in 6 mm simple well chambers with 100 μl E18 medium (Neurobasal Medium (Gibco) with 2% B27 (Gibco), 2.5% GlutaMAX (Gibco) and Antibiotic-Antimycotic). 8 µl cell suspension at 4 x10^6^/ml was loaded into each chamber of the microgroove device, and returned to cell culture incubator for 90 min. Then the device was topped up with E18 medium, using a multichannel pipette with two tips to load 100 μl per inlet pairwise to avoid flow across the chamber. Cells were maintained for 5-14 days without changing the medium, taking care to reduce medium evaporation. Cell growth and density were monitored using a simple custom designed upright microscope, dubbed ChipMONQ (Mono-Ocular Neuron Quantitator for Chip Microscopy), consisting of a 4X 0.13NA air objective (Nikon), tube lens (200 mm focal length, Thorlabs), ocular (Leica), flashlight for illumination, and a smartphone for image capture, visualisation, and digital zoom, see Figure S4.

#### Cell Staining

Primary neurons were stained with 1 μM SiR-Tubulin (Spirochrome Ltd., Stein am Rhein, Switzerland) and 10 μM verapamil (Spirochrome Ltd.), and where indicated 40 nM MitoTracker Orange CMTMRos (Thermo Fisher) in fresh E18 medium for 1 hour, before swapping to imaging medium (E18 medium with 20 mM HEPES pH 7.4). As shown in Figure 2, for the simple well chambers, a PDMS ‘fence’ 2 mm high was added around the chambers, then the chamber block was removed, and more imaging media was added, making sure that the sample remained moist at all times. For the microgroove chamber, the top layer was removed making sure not to disturb the thin bottom layer, and a cover slip was added to stop evaporation during imaging.

For immunofluorescence, primary neurons were fixed (2% paraformaldehyde, 2% glutaraldehyde, 100 mM cacodylate pH 7.4, 2 mM CaCl_2_) for one hour at room temperature and washed with PBS. Samples were incubated in blocking solution (PBS with 10% goat serum, 0.05% saponin) for one hour, stained with anti-VAMP2 (mouse monoclonal 104211, Synaptic Systems, Goettingen, Germany) and anti-calnexin (rabbit polyclonal ab22595, Abcam, Cambridge, UK) over night, washed, and goat anti-rabbit-Alexa Fluor 647 (Thermo Fisher) and goat anti-mouse-Alexa Fluor 488 (Thermo Fisher) for two hours, and washed. Samples were then stained with DiO (Vybrant, Thermo Fisher) and washed.

### Suppl. Note 7: Fibroblast Cultures

#### Fibroblast Cultures

Human Foreskin Fibroblasts from the American Type Culture Collection (ATCC) were cultured in DMEM with 10% FBS, 2.5% GlutaMAX and antibiotic-Antimycotic, in simple well chambers on waveguide chips.

#### Cell Staining

Fibroblasts were stained with 1 μM SiR-Actin (Spirochrome Ltd) and 10 μM verapamil for one hour. They were then fixed with 4 % PFA in PBS for 10 minutes before imaging.

### Suppl. Figures

**Figure S1:**
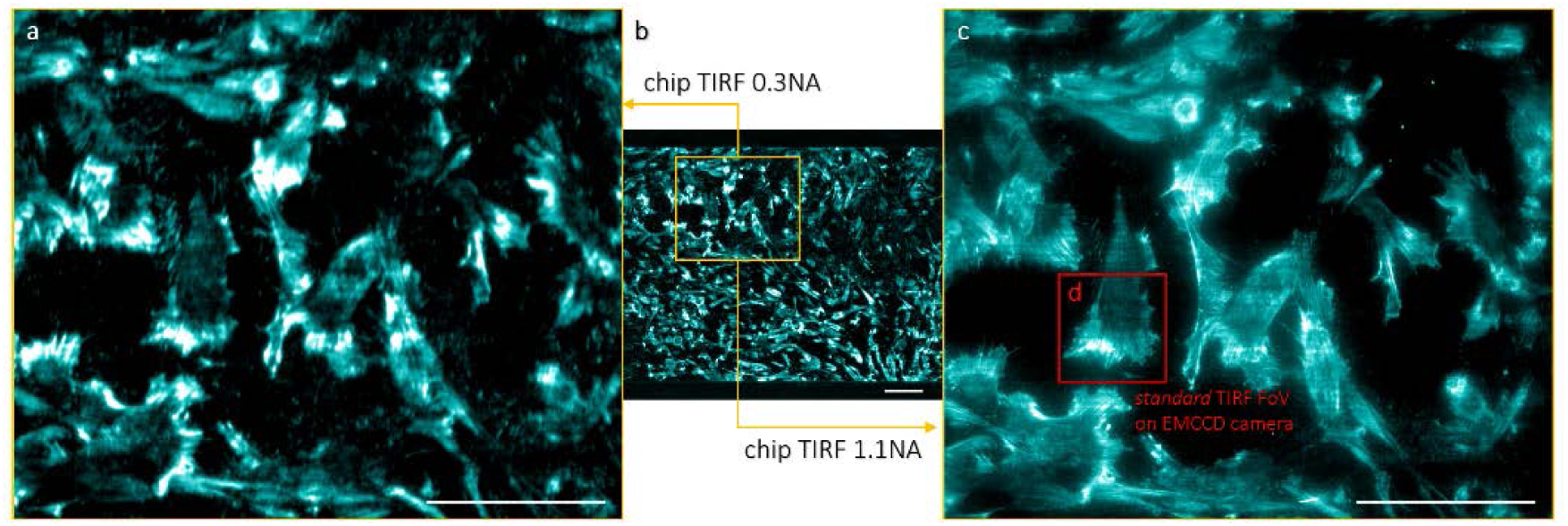
TIRF illumination provided by waveguides comes with the possibility of an almost arbitrarily large TIRF illumination area decoupled from the detection path. All panels show fixed fibroblasts on the same waveguide labelled with SiR-Actin. a) Zoom in on indicated region in b) which shows full width of the illuminated waveguide using a 0.3NA 10x water dipping objective. c) The same region as in a) but instead imaged using a 1.1NA 25x lens, providing higher magnification but in a smaller FOV. The red square (panel d), indicates the FOV provided using standard TIRF microscopy, i.e. 60 x 60 µm. The scale bars are 100 µm.

**Figure S2:**
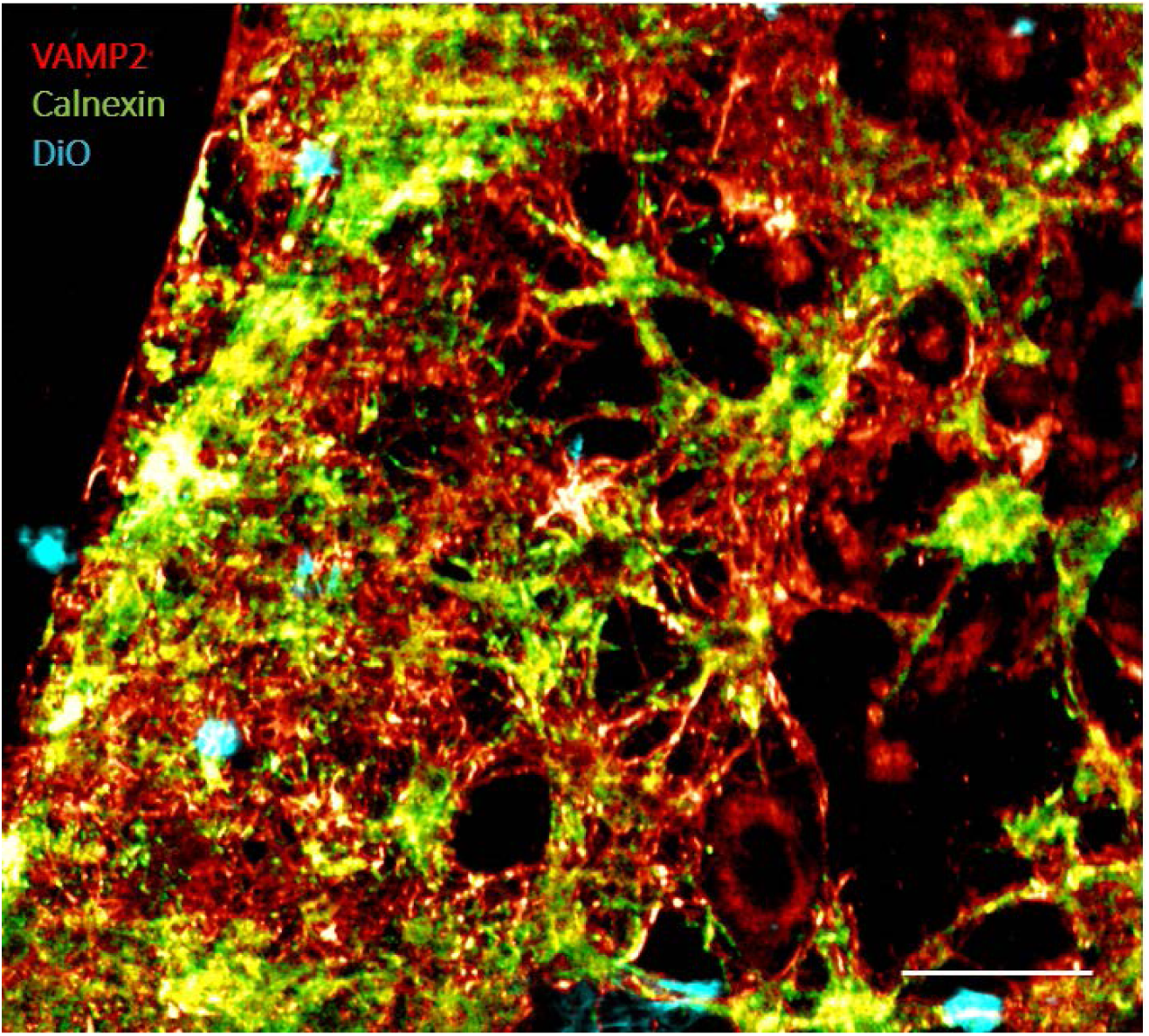
Via a suitable set of dichroic mirrors and emission filters together with three CMOS cameras synchronised to four laser lines, the ChipScope is equipped for simultaneous multi-colour TIRFM image acquisition. The image shows proof of concept three-colour waveguide TIRF imaging of rat hippocampal neurons immunolabelled for VAMP2 (red) and Calnexin (green) in addition to the membrane probe DiO (cyan). The scale bar is 100 µm.

**Figure S3:**
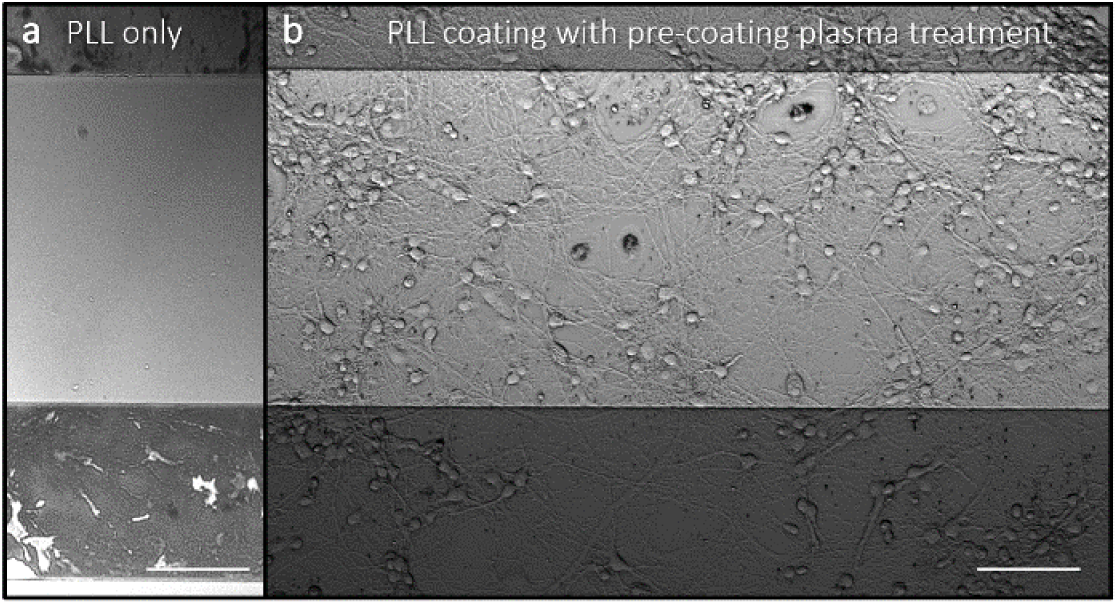
Cell culture on waveguides can require different surface treatment compared to what is used for standard microscopy glass coverslips. We found that countering waveguide hydrophobicity with low pressure oxygen plasma treatment right before poly-l-lysine coating greatly improved cell viability for both hippocampal and retinal cell cultures. a) Example of on-chip hippocampal cell culture without plasma surface treatment, b) example of on-chip hippocampal cell culture after pre-coating plasma treatment. The scale bars are 100 µm.

**Figure S4:**
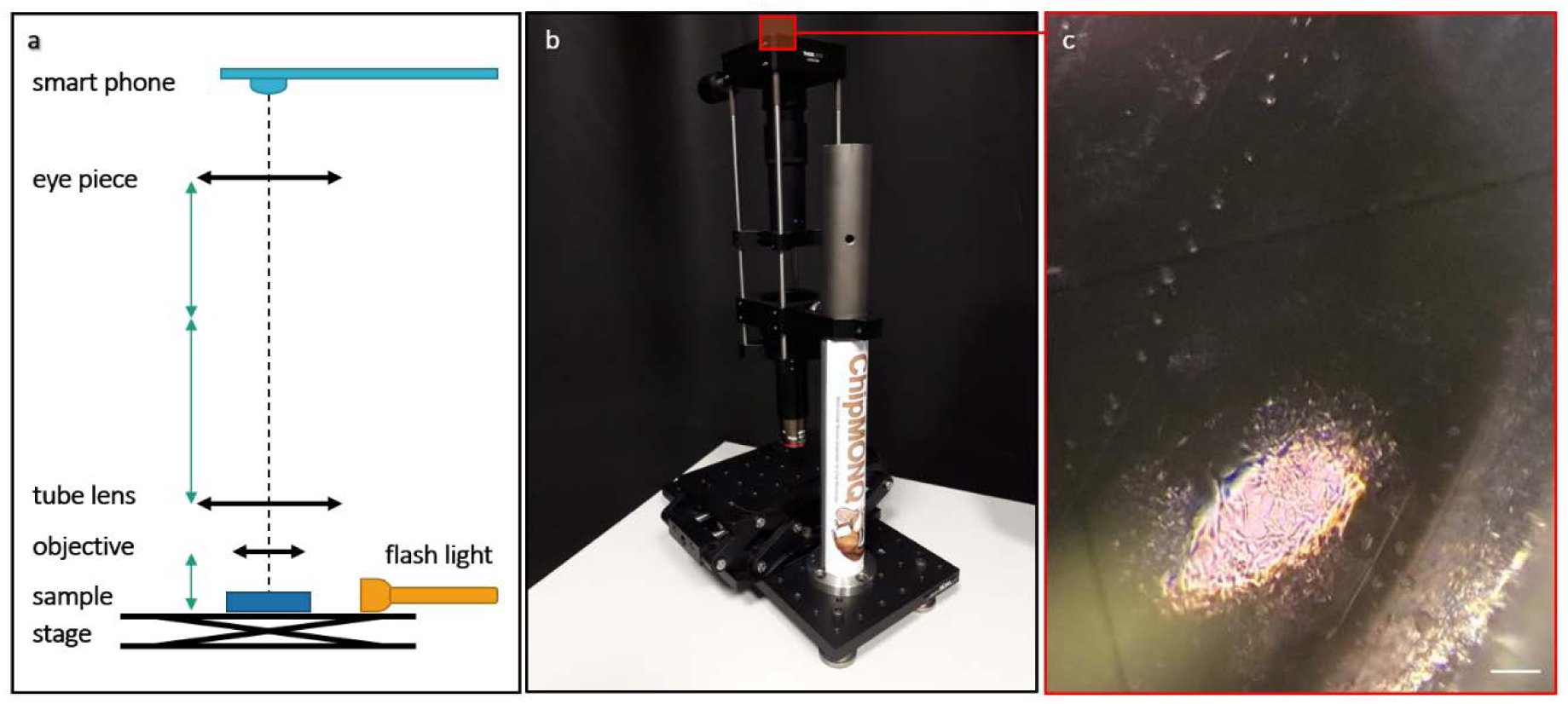
The custom-made Mono-Ocular Neuron Quantitator for Chip Microscopy (ChipMONQ) was used for monitoring cell growth and health during culturing before TIRFM on the ChipScope. As conventional cell culture microscopes are based on transmitted light, they are not applicable to samples on non-transparent substrates like silicon. To still be able to monitor cell growth and health on waveguides, we custom-built a simple upright microscope that could be easily moved and fitted on a cell lab bench. a) Sketch of the optical set-up. b) Picture of the ChipMONQ. c) Image of hippocampal cultures imaged using the ChipMONQ and a smartphone camera. The scale bar is about 200 µm.

**Figure S5:**
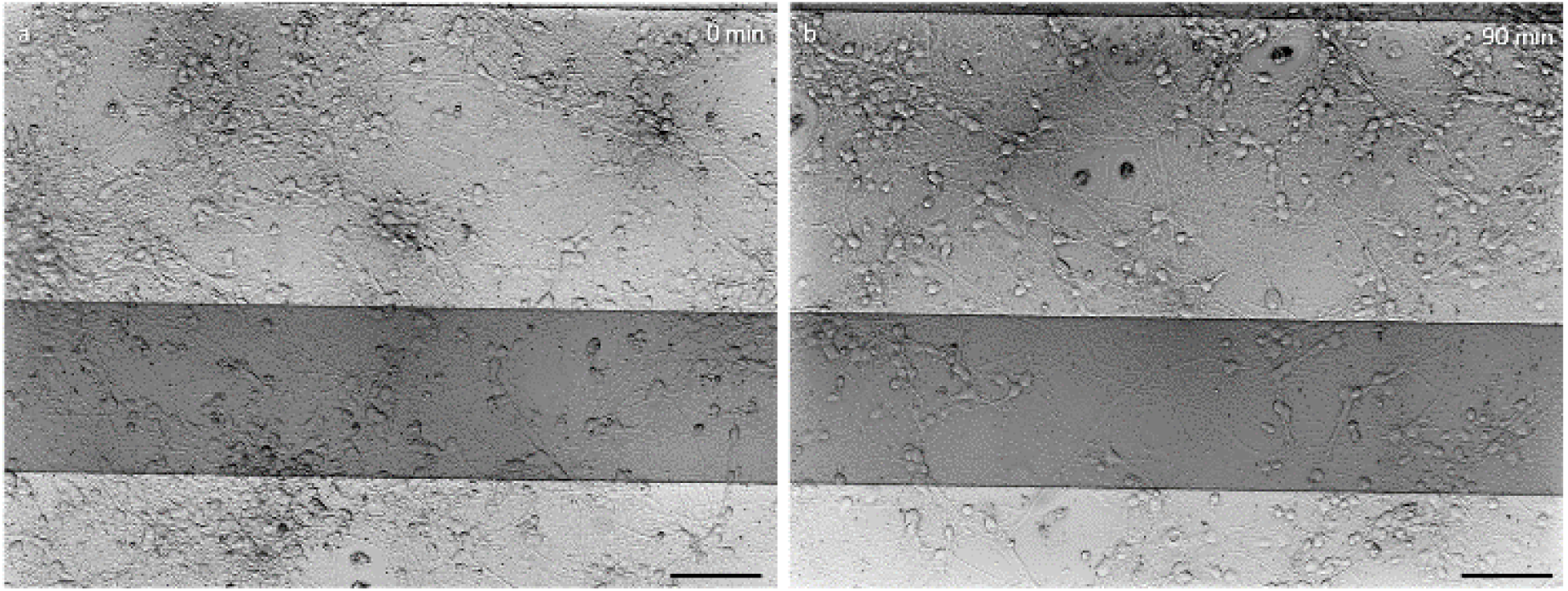
The ChipScope heating and environmental chamber enables long-term chip microscopy of delicate samples like mammalian hippocampal cell cultures. The images show brightfield overview images at the beginning (*left*) and after 90 min of imaging (*right*). Brightfield microscopy images were used since this is how neuronal cell cultures are usually monitored. The living neurons appear in healthy condition both before and after the prolonged period outside the conventional cell culture incubator, demonstrating the capabilities of the *ChipScope* incubator.

**Figure S6:**
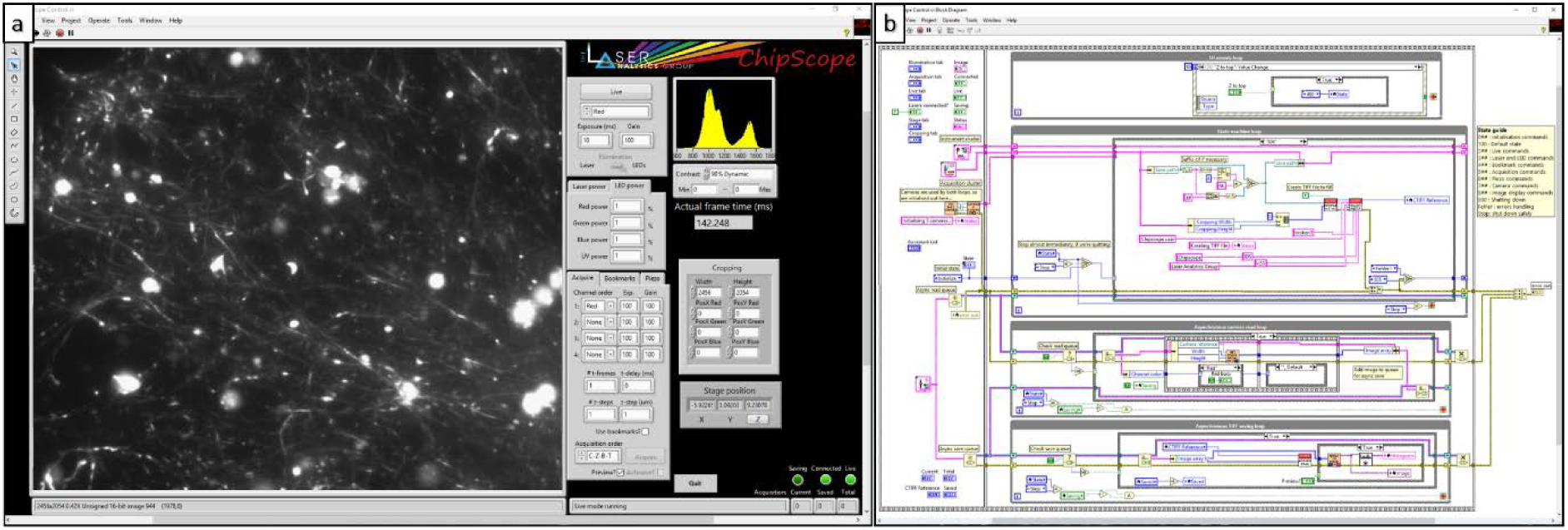
The ChipScope is controlled via a custom-made LabVIEW software. a) Front panel user interface, b) back panel block diagrams. A detailed description of the software is provided in Suppl. Note 2.

